# Spatially and temporally varying selection influence species boundaries in two sympatric *Mimulus*

**DOI:** 10.1101/2022.12.07.519524

**Authors:** Diana Tataru, Emma C. Wheeler, Kathleen G. Ferris

## Abstract

Spatially and temporally varying selection can maintain genetic variation within and between populations, but it is less known how these forces influence divergence between closely related species. We identify the interaction of temporal and spatial variation in selection and their role in either reinforcing or eroding divergence between two closely related *Mimulus* species. Using repeated reciprocal transplant experiments with advanced generation hybrids we compare the strength of selection on quantitative traits involved in adaptation and reproductive isolation in *Mimulus guttatus* and *Mimulus laciniatus* between two years with dramatically different water availability. We found strong divergent habitat mediated selection on traits in the direction of species differences during a drought in 2013, suggesting that spatially varying selection maintains species divergence. However, a relaxation in divergent selection on most traits in an unusually wet year (2019), including flowering time which is involved in pre-zygotic isolation, suggests that temporal variation in selection may weaken species differences. Therefore, we find evidence that temporally and spatially varying selection may have opposing roles in mediating species boundaries. Given our changing climate, future growing seasons are expected to be more similar to the dry year, suggesting that in this system climate change may actually increase species divergence.

## Introduction

The strength and direction of natural selection on a single population varies both over time and space. This spatial and temporal variation may either advance or erode population and species differences (Hereford 2009). Evolutionary biologists have long studied the phenotypic and genetic effects of varying natural selection (Darwin 1859). Spatially varying selection often leads to the maintenance of genetic variation both within and between populations (Mojica et al. 2012, Sheth & Angert 2016; Walter et al. 2020). Spatial environmental heterogeneity can maintain genetic variation within species through balancing selection mechanisms such as overdominance, GxE interactions, or frequency-dependent selection (reviewed in Delph and Kelly 2013). Measures of genetic variation at species’ range edges (Sheth and Angert 2016) and within different ecotypes (Walter et al. 2020) have experimentally confirmed the importance of the resulting increase in genetic variation in population survival and adaptation. Temporally varying selection can increase genetic variation in the short term (Borash et al. 1998; Mojica et al. 2012), but the maintenance of variation may be less stable due to more frequent changes in the direction of selection (reviewed in Kassen 2002; Wilson et al. 2006; Siepielski et al. 2009). The maintenance of genetic variation within a species is essential for long term persistence as it allows for local adaptation across environmental conditions (Barrett & Schluter 2007) and may improve species resilience by increasing the speed of evolutionary response to shifting environments due to climate change (Franks et al. 2007; Bell & Gonzalez 2009; Hoffman & Sgro 2011).

While spatially and temporally varying selection have been studied extensively in the context of local adaptation and genetic variation within species (Hereford et al. 2009; Mojica et al. 2012; review in Savolainen et al. 2013; Anderson et al. 2014; Anderson et al. 2021), it is less well known how spatially and temporally varying selection affects divergence and reproductive isolation between species. This is particularly important to understand when species occur sympatrically and are incompletely reproductively isolated. Adaptation to different environments can lead to and reinforce reproductive isolation between populations through the evolution of pre-zygotic (i.e. temporal, mating or habitat isolation) or extrinsic post-zygotic isolating barriers (i.e. reduced hybrid fitness), facilitating species divergence (Coyne & Orr 2004; Lowry et al. 2008; Mimura & Aitken 2010; Walter et al. 2020).

Differential adaptation is considered an important driver of speciation (Sobel et al. 2010). Recently diverged sympatric species are well suited for examining how fluctuating selection may affect the maintenance of reproductive isolation. To investigate how shifts in selection over time and space affect ecologically important and reproductively isolating traits we replicate a previously published reciprocal transplant (Ferris and Willis 2018) and identify patterns contributing to or eroding divergence between sympatric Monkeyflower species in two years with dramatically different snowpack levels.

Due to their sessile nature, plants often experience strong divergent selection in heterogeneous environments (Kalisz 1986), which makes them ideal for measuring natural selection on quantitative traits. Species with incomplete reproductive isolation can be easily interbred to break up within-species linkage disequilibrium and phenotypic correlations through multiple generations of recombination (Nagy 1997). With the creation of advanced generation interspecific hybrids, selection can be measured on individual traits because each trait is segregating in a randomized genetic background. Annual plants have short life cycles, allowing for experimental designs that can closely track life history traits, fitness, and generational differences. All of these factors make plant species ideal for studying spatial and temporal variation in selection in their native habitats. The *Mimulus guttatus* species-complex is excellent for answering questions about variation in environmental selection because it is a morphologically and ecologically diverse group of closely related species (Wu et al. 2008). The wide-ranging and largely outcrossing *M. guttatus* occupies continually moist seeps, while the highly self-fertilizing *M. laciniatus* occurs in granite outcrops throughout the Sierra Nevada Mountain Range in California. These granite outcrops exhibit shallow soils, ephemeral water supply from snowmelt, intensive light, and more extreme temperatures than nearby *M. guttatus* habitat (Ferris et al. 2014). A previous study found that *M. laciniatus* has several traits that are adaptive in its native rocky outcrops; small flowers, early flowering time, and lobed leaves (Ferris and Willis 2018).

Flowering time is known to be divergent in sympatry and allopatry between the two species, with *M. laciniatus* flowering earlier than *M. guttatus*, which confers temporal reproductive isolation (Ferris et al. 2017). Ferris and Willis (2018) also found selection against immigrants in each species’ native habitat, indicating habitat isolation. Post-zygotic isolation is not well studied in these species, however late generation hybrids have been observed in the wild (Vickery 1964; Ferris et al. 2014; Tataru personal observ), and are easily created in the lab suggesting that intrinsic barriers are not significant. Therefore while temporal and habitat isolation form pre-zygotic barriers between the two species, they are incompletely reproductively isolated. Flower size and flowering time are known as “magic traits”, or adaptive traits that also contribute to reproductive isolation and can facilitate or maintain species divergence in the face of gene flow (Servedio et al. 2011). While not directly associated with reproductive isolation, *M. laciniatus*’ distinctive lobed leaf shape is also adaptive in its environment and thus could contribute to habitat isolation between the species (Bright & Rausher 2008; Ferris & Willis 2018). Quantifying differential selection in native species’ specific sites of *M. guttatus* and *M. laciniatus* allows us to investigate the stability of divergence in traits involved in both temporal and habitat isolation.

To examine temporal and spatial variation in selection, we replicated a reciprocal transplant experiment originally conducted during the historic California drought of 2013 (Ferris and Willis 2018) in the summer of 2019, an unusually high snowpack year in the Sierra Nevada (cdec.water.ca.gov; Tuolumne River Basin Snowpack Average Pct. Of April 1, 2013: 52%, 2019:176%). In this study we aim to answer the following questions: 1) What are the patterns of temporal and spatial variation in selection on quantitative traits involved in local adaptation and reproductive isolation within these two species’ different habitats? 2) How do those patterns potentially reinforce or erode divergence between the two *Mimulus* species? The 2013 experiment found divergent selection on flowering time, plant height, and leaf shape between the species’ habitats in the direction of species differences (Ferris & Willis 2018).

Differential seasonal soil moisture patterns between habitats were strongly associated with local adaption in the experiment, supporting prior research within the *Mimulus guttatus* species complex (Hall & Willis 2006; Kooyers et al. 2015; Mantel & Sweigart 2019). To understand temporal variation in divergent selection between the species we repeated this original reciprocal transplant in 2019 with the same fourth generation *M. guttatus* and *M. laciniatus* hybrid population. We predicted there would be spatially varying selection in the direction of species differences which would then reinforce species divergence and habitat isolation, maintaining species boundaries over time. We also expected that the strength of selection would vary from a drought season (2013) to a very wet season (2019). Given the effects of global climate change California’s drought is only expected to increase in severity over the coming decades. Therefore selection patterns similar to those in the drier transplant year are expected to be more common in the future. Understanding the effect of rare wet years on patterns and strengths of selection will give us further insight into the constraints on drought adaptation and processes of divergence.

## Methods

### Reciprocal Transplant Design

We conducted our repeated reciprocal transplant from beginning of June to October 2019 in Yosemite National Park, CA, USA. We replicated an experiment performed in 2013 by Ferris and Willis (2018), using the same four sites and seeds from the same F_4_ hybrid population. The four sites were as follows: two undisturbed granite outcrops with native *M. laciniatus* growing on moss, Olmstead Point (Granite 1, 8500 ft) and Yosemite Creek (Granite 2, 7500 ft; Figure 1), and two undisturbed meadows with native *M. guttatus* growing near a standing seep, Little Meadow (Meadow 1, 6200 ft) and Crane Flat (Meadow 2, 6000 ft; Figure 1). These sites were chosen to maximize the similarity between the developmental stages of transplanted and native plants of each species (Ferris & Willis 2018).

**Figure 1.**
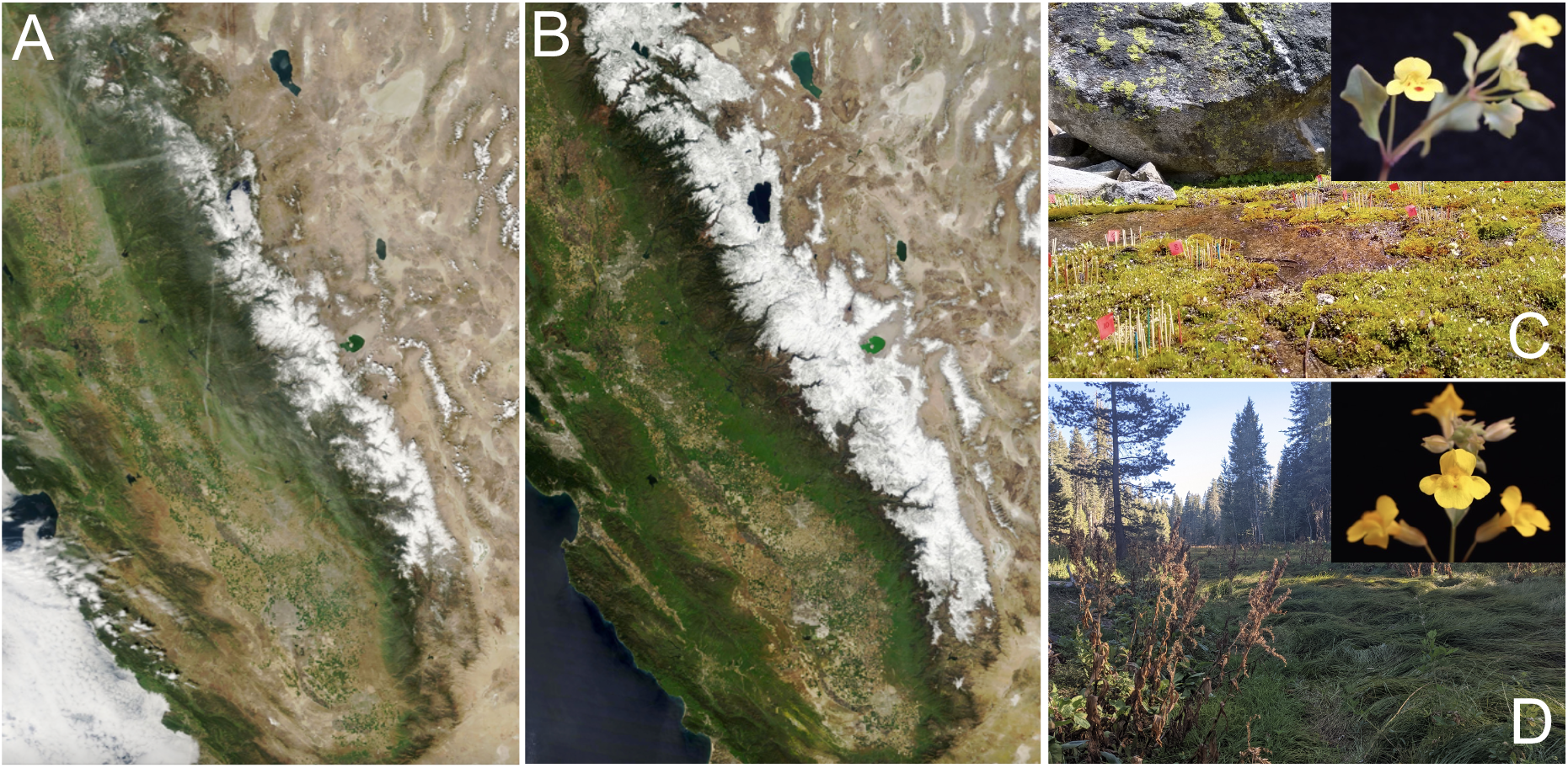
A) Imagery of Sierra Nevada snowpack March 29, 2013 and B) March 31, 2019 from the NASA MODIS Terra Satellite, courtesy NASA NSIDC. C) Granite Outcrop 2 experimental set-up with a photo of native *Mimulus laciniatus*. D) Meadow 2 with a photo of native *Mimulus guttatus*.

The experimental F_4_ hybrids used in both the 2013 and 2019 experiment were bred in 2013 in the Duke University greenhouse using inbred *M. guttatus* (YVO 6) and *M. laciniatus* (WLF 47) lines (visualization of crossing design in Ferris and Willis 2018). F_3_ hybrids were created by crossing the parental inbred lines to generate F_1_ hybrids which were self-fertilized to generate F_2_s, followed by randomly intercrossing one generation of 200 different maternal hybrid lines (Ferris & Willis 2018). We pooled 30 seeds from 200 maternal F_3_s to make an outbred, pooled F_4_ population. We cold-stratified parental and hybrid seeds at 4°C for 10 days, and then germinated plants for one week in growth chambers at UC Berkeley. Due to low initial germination in 2019 we also germinated a second set of plants two weeks later in identical conditions in UC Merced growth chambers. At the cotyledon stage we transplanted seedlings one inch apart into 50 randomized blocks of 19 plants each (2 WLF47, 2 YVO6, 15 F_4_) at each of four field sites. We placed blocks 6-18 inches apart with toothpicks marking individual seedlings. Native *Mimulus* in each block and within one inch of the block were removed. Genotypes within blocks were randomized to account for neighbor effects, edge effects, and aspect. Due to limitations with germination and the timing of site access, the number of experimental plants at each site varied (Meadow 1: 921 total, 750 F_4_s, 100 *M. guttatus*, 71 *M. laciniatus;* Meadow 2: 900 total, 750 F_4_s, 100 *M. guttatus*, 50 *M. laciniatus*; Granite 1: 645 total, 600 F_4_s, 60 *M. guttatus*, 60 *M. laciniatus*; Granite 2: 792 total, 660 F_4_s, 88 *M. guttatus*, 44 *M. laciniatus*). Low sample sizes for parental species in 2019 were due to limited germination of seeds in greenhouses. Seedlings that died within a week were replaced so that transplant shock did not skew survival data.

### Environmental Measurements

To assess variation in potential environmental agents of selection over time and space, we took fine scale measurements of soil moisture, surface temperature, and light measurements at each site every week in 2019. We also measured the presence of herbivore damage on mature plants. Measurements were taken at half of the blocks in each site (25) at the same location within each block. Environmental measurements were selected to span micro-habitat variation in each site. In the 2013 experiment only soil moisture was measured. We analyzed environmental data from 2019 and re-analyzed 2013 data using linear mixed effects models in the nlme package in R (Pinheiro et al. 2022) to find associations between environmental measurements and plant survival. We used plant survival as a metric of fitness because it provides the most fine-scale temporal data. Survival was measured as the proportion of plants surviving in each block at the time of each environmental survey. We ran these models with data from each year both separately and grouped to identify specific selective forces within years as well as broader patterns across years. We grouped sites by habitat. To identify whether patterns in seasonal soil moisture decrease were comparable across years in each habitat, we ran linear mixed effects models with years combined, survival as the dependent variable and soil moisture, year, and date as fixed independent variables, and block as a random effect. We also ran linear mixed effects models with our 2019 data separately with survival as the dependent variable and soil moisture, habitat, time, soil surface temperature (2019) and light levels (2019) as fixed independent variables, and block as a random effect. In all models we tested for interactions between independent variables and determined the best fit model using AIC model selection using the R package MuMin (Barton 2009).

To understand the role and presence of herbivory across sites and years, we noted herbivory in the field on each individual as presence or absence. In each year herbivory data was taken on all individuals that flowered, but only F_4_ hybrids were analyzed because of insufficient sample size in the parents. To analyze whether herbivory and habitat are independent variables, we ran chi-squared tests to test for correlation between variables, and then created contingency tables to quantify differences. To understand the importance of herbivory in plant fitness, we ran ANOVA models using F_4_ fruit data as the dependent variable and herbivory presence/absence across years, habitat, and block as independent variables, as well as interactions between them. We used model selection with MuMin (Barton 2009) to find the best fit model.

### Phenotypic Measurements

To understand how selection was acting on plant phenology, mating system, and morphology in each species’ native habitat we surveyed plants every other day for survival and phenotypic measurements. On the day of first flower we measured the flowering time, plant height (soil to plant meristem), stigma-anther separation, corolla size (width and length), and leaf measurements. We took measurements with digital calipers. Stigma-anther separation was calculated by subtracting the length of the longest anther from the stigma length and is used as a relative metric of outcrossing. Stigma-anther separation was only measured in 2019. We collected the first true leaves once plants began to senesce but before leaves browned, and later measured leaf area and lobing index by digital scanning and analysis in the program Image J as described in Ferris et al. (2015). Briefly, leaf lobing is calculated as the convex hull area minus the true leaf area divided by convex hull area. Flower width and length were strongly correlated in both years and thus flower width is used in all subsequent analyses as our metric of flower size.

Once plants senesced we counted the number of fruits and collected individuals in labeled coin envelopes. In a lab, we counted seed number by pouring packets onto a grid and visually counting seeds. Seeds were then recounted by a different person to ensure accuracy. Fruit and seed numbers were used as our metrics of fitness. Correlation between seed and fruit number was calculated using Kendall’s correlation analysis.

### Phenotypic Selection Analysis

To understand linkage between traits and confirm estimates of selection (Mitchell-Olds and Shaw 1987), we ran a phenotypic trait correlation matrix for 2019 hybrid traits in separate habitats using the R package corrplot (Wei & Simko 2021). We report r as the correlation coefficient. A high positive or negative r value indicates strong correlation between traits, suggesting linkage between traits (Falconer & Mackay 1996). We compared trait means in hybrids and parents using t-tests to understand putative plasticity across environments and calculate trait variance (C_v_) by dividing the standard deviation of the traits by the reported means (Pelabon et al. 2020).

We analyzed phenotypic and fitness data of hybrids from 2019 to identify differences in the strength or direction of selection on phenotypes between years. The 2013 data used in the following analyses is published in Ferris and Willis (2018) and is re-analyzed with previously unpublished data from 2019. We used multivariate linear and quadratic phenotypic selection analyses in hybrids to quantify selection on quantitative traits standardized to a mean of 0 and SD of 1 (Lande & Arnold 1983; Mitchell-Olds & Shaw 1987), and using block nested in site as a random effect. We re-ran all multivariate analyses on all 2013 and 2019 trait and fruit data, and 2019 seed and trait data (S2). We separated the analysis into negative binomial and truncated poisson models to account for overdispersion of zeros in viable fruit and seed number. Similarly to Ferris and Willis (2018), we ran these models separately in each habitat to identify differential habitat selection. Only plants that survived to flowering were used in these analyses because all phenotypic data was collected on the day of first flower. The negative binomial model assessed whether a trait is associated with the production of any fruits, while the truncated poisson measured whether each trait is associated with the number of fruits produced. In 2013 we only collected fruit number data for our fitness metric, while in 2019 we collected both fruit and seed numbers. We repeated the above phenotypic selection analyses with 2019 seed data to identify differences in patterns of selection when fruits vs. seeds are used as a fitness proxy. We ran analyses using the R packages glmmadmb (Fournier et al. 2012) and glmmTMB (Brooks et al. 2007), and then conducted AIC model selection using the R package MuMin (Barton 2009). We report selection gradients (*β*) for all traits in our top models.

## Results

### Higher Fitness in a Wetter Year (2019)

We found higher hybrid fitness due to increased fecundity in 2019 than 2013, as we predicted with increased snow melt (Figure 4). Survival in 2019 was as follows; in Granite 1 no plants survived to flowering while at Granite 2 31% hybrids and no parents survived to flowering. In Meadow 1, we saw high hybrid (35%) and *M. guttatus* (18%) survival to flowering, but low survival for *M. laciniatus* (1%). In Meadow 2 we saw low survival to flowering across genotypic categories (8% hybrid, 8% *M. laciniatus*, 6% *M. guttatus*). Across all sites in 2019, 530 total F_4_ hybrids survived to flowering, a comparable sample size to the 525 hybrids which survived in 2013. Mean and total fruit number for hybrids in 2019 (Meadow: total fruit=521, mean=1.612; Granite total fruit=442, mean=2.167) was higher than in 2013 (Figure 4; Meadow: total fruit=18, mean=0.06; Granite: total fruit=285, mean=0.188). Survival and seed number in parental species suggested that in their native meadow habitat, *M. guttatus* individuals were more than twice as successful in surviving to reproduction (20 *M guttatus*; 9 *M. laciniatus*) and producing seeds (10 *M guttatus*; 5 *M. laciniatus*) than *M. laciniatus* (Figure S1). This differs from the dry 2013 transplant where there was only a slight advantage for *M. guttatus* in fecundity in its native meadow habitat, but a significant advantage in fecundity and survival for *M. laciniatus* in its native granite habitat (Ferris & Willis 2018). However, we are not able to identify fitness trade-offs and conduct a complete comparison between habitats in parental species lines in 2019 due to extremely low parental survival to flowering in granite (0 *M. laciniatus*; 1 *M. guttatus*). Low survival in our Granite 1 site was likely due to a logistical issue with planting time due to low initial germination and not patterns in local adaptation. Because we did not have enough plants germinate initially, we had to delay planting out the Granite 1 site by two weeks. This had a large impact upon fitness which was confirmed by a significant statistical difference in plant survival based on planting time (Figure S2; p<,.0001, F=77.2). Even with these logistical planting challenges we still saw significantly higher hybrid fitness in the 2019 transplant.

### Soil Moisture is Important for Plant Survival Across Time and Space

We found a significant association between seasonal trends in soil moisture and plant survival across all years and habitats, while light intensity and surface temperature were not significantly associated with survival in 2019 (Table S1). There was a significant difference in soil moisture levels between habitat types in 2013 but not 2019 (2013: p<.0001, df=1922, F=101.746; 2019: p=0.138, df=719, F=2.2053), and significant interactions between soil moisture, time, and habitat. These interactions indicate that there is a significant difference in seasonal soil moisture decrease between habitats. Across years, granite sites had a plateau of high early season soil moisture with a steep drop midway through the season, while in meadows the decrease in soil moisture over time was shallow and mostly linear (Figure 2). Absolute levels of soil moisture were 2-3X higher in 2019 than 2013. Analyzing data from each habitat separately, soil moisture, time, site, and year were all significantly associated with survival, however the best model for meadows included an interaction between time and site, while the best model for granite included an interaction for soil moisture and time. These interactions suggest that meadow sites had differential survival over time based on site, while survival in granite outcrops was strongly associated with soil moisture decrease over time. The lack of interaction of year with soil moisture suggests that while there was significantly higher absolute soil moisture in 2019, there are comparable patterns in seasonal soil moisture decrease across years.

**Figure 2.**
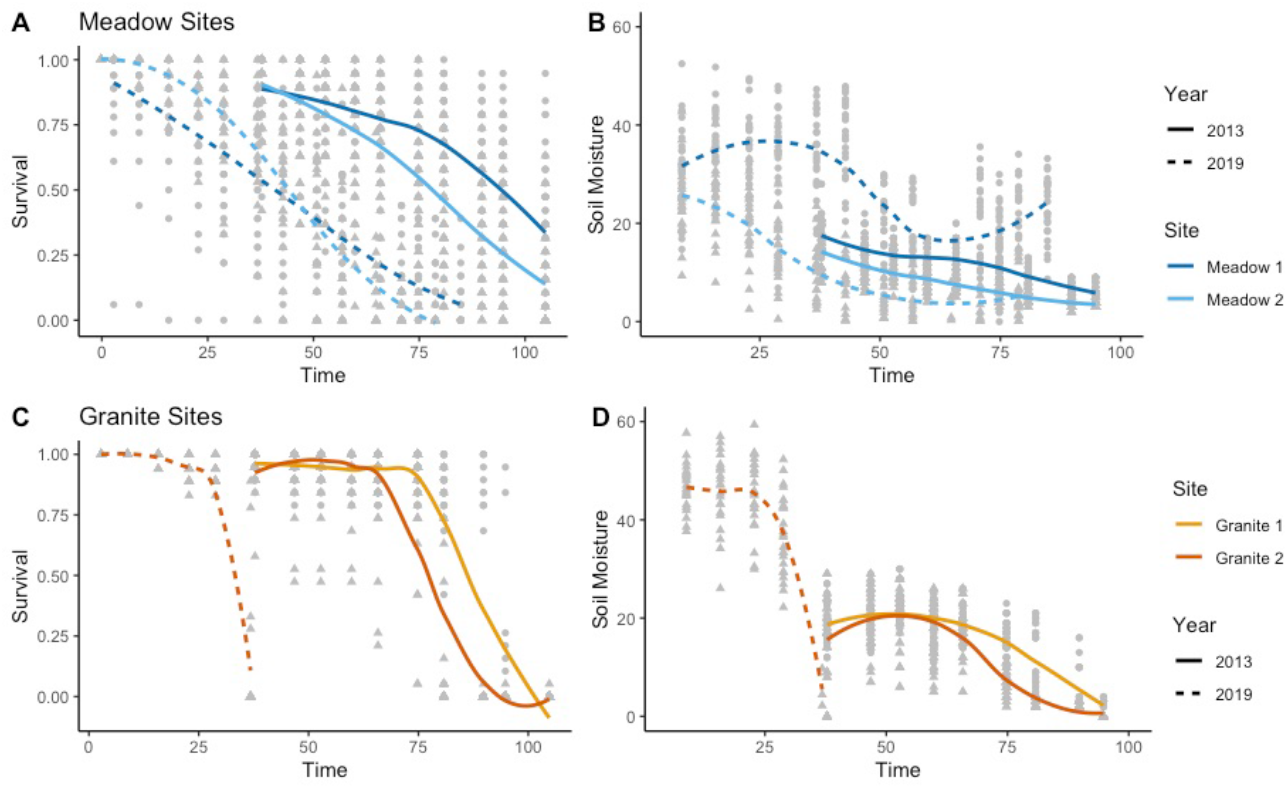
Survival since planting day and soil moisture in %volumetric soil moisture in both years of the reciprocal transplant. Meadow sites are indicated with shades of blue and granite sites are indicated with shades of orange. 2013 data is published in Ferris and Willis (2018) and re-analyzed here.

### Herbivory Differs Across Species’ Habitats

We found that herbivory pressure differed significantly between habitats in both 2013 (X2=62.868, p<.0001) and 2019 (X2=19.584, p<.0001), with a stronger relationship between herbivory and habitat in 2013 than 2019 (Figure S4). Contingency tables of habitat type and herbivory showed that there was a higher proportion of herbivory in meadows than granite outcrops in both years, with a higher proportion of herbivory in meadows in 2013 than 2019 (2013 meadow: 14.25% granite: 2%; 2019 meadow: 7% granite: 2.27%). The best fit ANOVA model of the effect of herbivory on fitness (fruit production) in 2013 included herbivory, site, and an interaction between herbivory and site, however only site was significant (F=24.1, p<.0001, df=3). The best fit ANOVA model looking at the effect of herbivory on fruit production in 2019 included herbivory, site, and block. Therefore, herbivory did differ between habitats in both years and affected fruit set.

### Phenotypes are Expressed Differently Across Habitats

While phenotypes are expressed differently across habitats, trait expression is also affected by strong correlations between traits. Differential trait means and coefficients of variation (C_v_) indicate significant variation in leaf size, leaf lobing, and flowering time in experimental hybrids across habitats and between years (Table S2). In both years, granite outcrop hybrids had earlier mean flowering time and wider flowers than those grown in meadows. In the 2019 phenotypic trait correlation matrix (Figure S5) for meadow hybrids, we found strong positive correlations between flower width and plant height, flower width and flowering time, and plant height and flowering time (r>0.6). In granite outcrops, we found positive correlations between flower width and plant height (r=0.39), flowering time and plant height (r=0.2), and leaf area and flower width (r=0.42). Similar to the 2013 experiment, we found a slight positive correlation between leaf shape and area (r<0.15). In 2013, flowering time was uncorrelated or weakly correlated with all traits across habitats (Ferris and Willis 2018), which was different from our 2019 findings. This suggests that unlike flower width and plant height, which show strong covariation across habitats and years, flowering time covariation with morphological traits is dependent on temporal environmental variation.

### Spatially Varying Selection Increases Species Divergence

Multilinear selection analyses of fitness in F_4_ hybrids indicated that our focal traits are under divergent selection between meadows and granite outcrops largely in the direction of species divergence across years. In 2019 we collected both fruit and seed number as fitness metrics, and found a significant correlation between fruit and seed number in both meadows (Figure S3; P<.001, correlation coefficient = 0.309, n = 1902) and the granite outcrop (S4; P=0.0036, correlation coefficient = 0.172, n=838). We first describe the results of the multivariate negative binomial followed by the zero-truncated poisson analysis, and focus on directional selection gradients because they account for trait correlations (Lande & Arnold 1983). The significant interaction between leaf shape and plant height (Meadow β=0.301, Granite β=-1.862), and negative interaction between flowering time and flower width (Meadow β=-0.486, Granite β=-1.612) in both habitats, suggest non-independence of selection on some traits (Lande & Arnold 1983, Mitchell-Olds & Shaw 1987). In the 2019 negative binomial analysis of whether a plant produced any viable fruits or not (Table 1), strength of selection on flower width and plant height is in the direction of species differences as predicted. However, not in the expected direction of species differences, the granite outcrop had stronger selection on later flowering time and rounder leaves than meadows (Figure 3). Using seeds as a metric of fitness in 2019, flowering time changed direction in selection from later flowering time in fruits to earlier in seeds in both habitats. As predicted by species differences, seed fitness indicated stronger selection for early flowering time in the granite outcrop.

**Table 1.** Negative Binomial analysis on whether or not a plant produced fruits or seeds (2013: n=525, 2019: n=530). Values indicate the selection gradient (*β*), or strength of selection. Significance of selection in a trait is indicated by “*” and significance codes are as follows: P-value < 0.001***; 0.01**; 0.05*. Missing values (dashes) indicate traits were not included in the best fit model for the year/habitat combination. 2013 data is published in Ferris and Willis (2018) and re-analyzed here.

| Trait | Meadow |  |  | Granite |  |  |
| --- | --- | --- | --- | --- | --- | --- |
| Year | $\beta$ 2013 (fruits) | $\beta$ 2019 (fruits) | $\beta$ 2019 (seeds) | $\beta$ 2013 (fruits) | $\beta$ 2019 (fruits) | $\beta$ 2019 (seeds) |
| Leaf Lobing (L) | -0.273 | -0.332 | - | 0.066 | -1.104* | - |
| Flower Width (FW) | -0.279 | 0.255 | 0.773** | 0.331* | -1.886* | - |
| Plant Height (PH) | 1.329* | -1.408*** | 0.464 | -1.256 | -1.384*** | - |
| Flowering Time (FT) | -0.216 | 1.289*** | -0.333 | -0.957*** | 2.444** | -0.987 |
| L*PH | - | 0.301 | - | - | -1.862*** | - |
| FT*FW | - | -0.486 | - | - | -1.612* | - |
| PH*FT | - | - | -0.519 | - | - | - |

**Figure 3.**
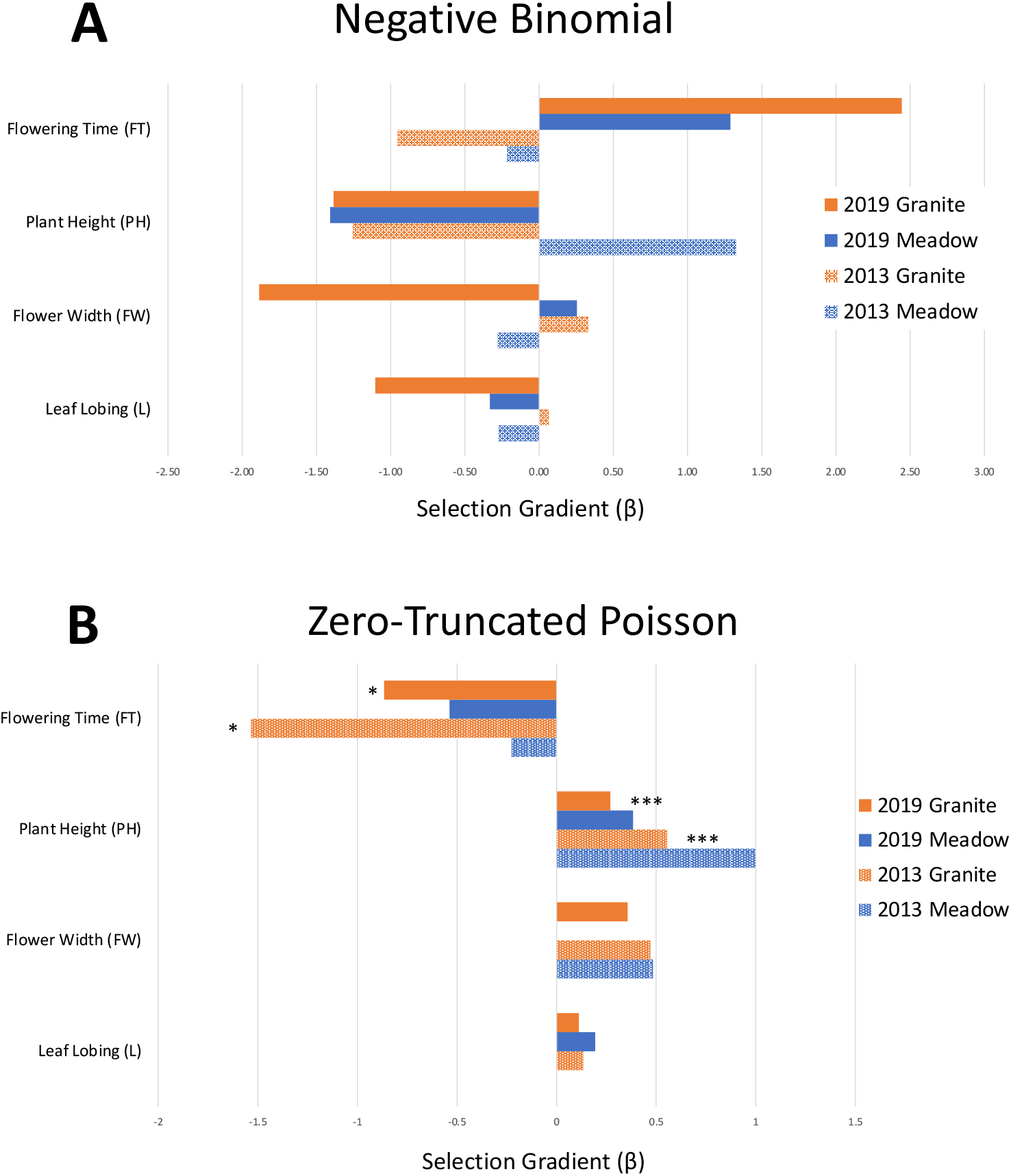
Visual representation of selection gradients (ß) based on fruit number from A.) Negative Binomial Models and B.) Zero-Truncated Poisson Models for 2013 (blue) and 2019 (orange). * indicate significant differences across years. 2013 data is published in Ferris and Willis (2018) and re-analyzed here.

The truncated poisson analysis using fruit number as the fitness metric (Table 2) in the 2019 granite outcrop showed similar results to 2013 in the direction of species differences, with stronger selection on early flowering time in the granite outcrop than meadows, and stronger selection for taller plants in meadows than granite outcrop. Truncated poisson models using seeds as the metric for fitness showed largely similar directions but differing strengths in selection as models using fruit number (Table 2). Similar to the binomial model, flowering time switches direction in selection when using fruits and seeds as fitness metrics. This switch, along with stronger selection for taller plants in the granite outcrop than meadows, does not track predicted species differences as closely as selection gradients based on fruits. We found no significant quadratic selection gradients.

**Table 2.**
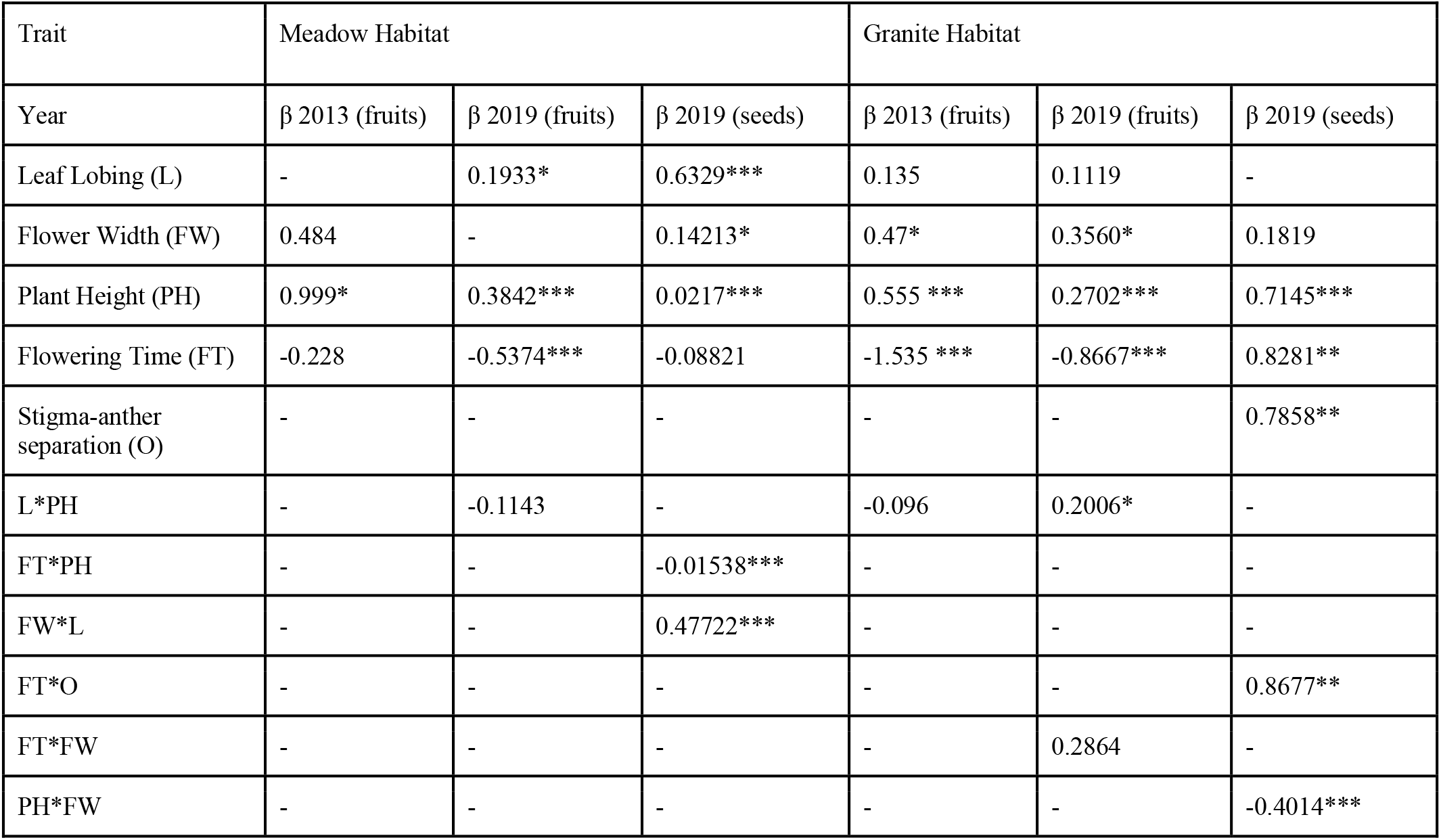
Zero-truncated poisson analysis on number of fruits or seeds produced if any were produced (2013: n=525, 2019: n=530). Values indicate the selection gradient (*β*), or strength of selection. Significance of selection in a trait is indicated by “*” and significance codes are as follows: P-value < 0.001***; 0.01**; 0.05*. Missing values (dashes) indicate traits were not included in the best fit model for the year/habitat combination. 2013 data is published in Ferris and Willis (2018) and re-analyzed here.

| Trait | Meadow Habitat |  |  | Granite Habitat |  |  |
| --- | --- | --- | --- | --- | --- | --- |
| Year | $\beta$ 2013 (fruits) | $\beta$ 2019 (fruits) | $\beta$ 2019 (seeds) | $\beta$ 2013 (fruits) | $\beta$ 2019 (fruits) | $\beta$ 2019 (seeds) |
| Leaf Lobing (L) | - | 0.1933* | 0.6329*** | 0.135 | 0.1119 | - |
| Flower Width (FW) | 0.484 | - | 0.14213* | 0.47* | 0.3560* | 0.1819 |
| Plant Height (PH) | 0.999* | 0.3842*** | 0.0217*** | 0.555 *** | 0.2702*** | 0.7145*** |
| Flowering Time (FT) | -0.228 | -0.5374*** | -0.08821 | -1.535 *** | -0.8667*** | 0.8281** |
| Stigma-anther separation (O) | - | - | - | - | - | 0.7858** |
| L*PH | - | -0.1143 | - | -0.096 | 0.2006* | - |
| FT*PH | - | - | -0.01538*** | - | - | - |
| FW*L | - | - | 0.47722*** | - | - | - |
| FT*O | - | - | - | - | - | 0.8677** |
| FT*FW | - | - | - | - | 0.2864 | - |
| PH*FW | - | - | - | - | - | -0.4014*** |

### Strength of Selection Varied Across Years

While direction of selection was largely the same across years, we did find weaker selection (Figure 3) and higher average fruit production in 2019 (Figure 4), supporting our hypothesis that increased water availability should relax selective pressures in 2019. To detect temporal variation in selection we compared the results of our phenotypic selection analyses between our 2013 and 2019 field experiments. Combining data cross years and quantifying the interaction between year and traits using fruit number as our fitness metric, we found significant interactions between year and flowering time (β=0.6022, p=0.0239) and year and plant height (β=-0.39244, p=0.000285) in the granite habitat truncated poisson analysis. Therefore selection on both flowering time and height differed significantly across years (Figure 3). In the negative binomial selection analysis we found a change in direction of selection across years in both habitats. In meadows, the direction of selection changed from tall plants and early flowering (2013) to strong selection for short plants and late flowering (2019). In the granite outcrop, the direction of selection changed from more lobed leaves, larger flowers, and early flowering (2013), to strong selection for less lobing, smaller flowers, and later flowering. Overall temporal variation in direction of selection on traits involved in adaptive species divergence such as flowering time and flower size shows weakened divergent selection in the direction of species differences in 2019. In the truncated poisson selection analysis there were no changes in the direction of selection between years, but strength of selection was weaker in 2019 on traits such as plant height and flower size. In 2019, strength of selection for early flowering time was stronger in meadows, but weaker in granite outcrops than in 2013 which is in the opposite direction of species divergence. Overall, this indicates that the strength of divergent selection between species native habitats was weaker in 2019 than 2013.

**Figure 4.**
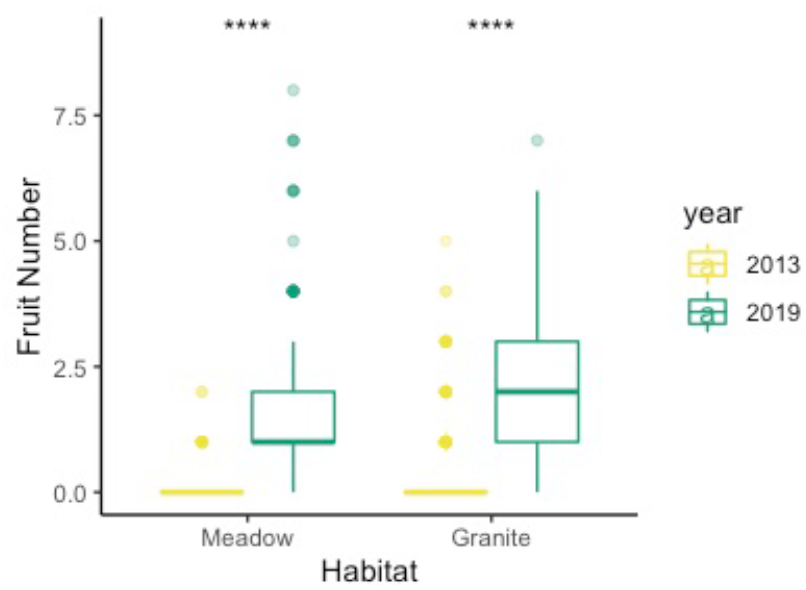
Total fruit number across years and habitats (2013 Meadow: total fruit=18, mean=0.06; 2019 Meadow: total fruit=521, mean=1.612; 2013 Granite: total fruit=285, mean=0.188; 2019 Granite total fruit=442, mean=2.167). Yellow boxes are 2013 data and green boxes are 2019 data. **** indicate significant differences in fruit number between years (P<.0001). 2013 data is published in Ferris and Willis (2018) and re-analyzed here.

## Discussion

Spatially and temporally varying natural selection are known to be important factors in the maintenance of genetic variation within species (Charlesworth 2015; Sheth & Angert 2016), however few studies have explored how fluctuating selection affects divergence and reproductive isolation between species adapted to different local habitats (but see Grant & Grant 2020, Anderson et al. 2015). In our repeated reciprocal transplant experiment in two closely related sympatric *Mimulus* species, we found that temporal differences in abiotic conditions shift the strength of spatially divergent selection. We observed selection for increased trait divergence and local adaptation under drought, but selection for decreased trait divergence in a year with high water availability.

These patterns presented differently depending on models, through either shifts in direction of selection in the binomial model or shifts in selection strength in the truncated poisson model. Both cases support decreased divergent selection in 2019 on traits involved in local adaptation and reproductive isolation between *M. laciniatus* and *M. guttatus*. Spatially varying selection across years remains mostly in the direction of species differences, indicating that local adaptation, or habitat isolation, plays an important role in maintaining divergence between the two sympatric species. However, in years with more relaxed divergent selection between habitats we may see a weakening of species differences. This is particularly important in traits that play a role in reproductive isolation between *M. laciniatus* and *M. guttatus* such as flowering time and mating system. The relaxation of divergent selection that we observed on flowering time in the wetter 2019 field season could lead to higher levels of interspecific gene flow in these incompletely reproductively isolated species and if frequent, could eventually lead to species fusion (Coyne & Orr 2004). However, given that 2019 was a rare snow pack year and long term trends in Sierra Nevada snow pack show persisting patterns of decreased snow, it is a much more likely outcome that the species will be under increased divergent selection in the future. Measuring spatial and temporal variability in selective pressures on these two sympatric *Mimulus* gives us insight into how species persist in harsh habitats, and how species’ divergence may change over time, an especially crucial topic of study in the current age of accelerated climate change.

### Differences in seasonal soil moisture between habitats may drive divergent selection

While local adaptation has been documented in many plant species, the environmental factors driving that adaptation are poorly understood in most cases (Kawecki & Ebert 2004; Hereford et al. 2009). We found evidence for strong temporally and spatially varying selection correlated with differences in soil moisture across years and different habitats of *M. guttatus* (meadows) and *M. laciniatus* (granite outcrops). We predicted that heavy snowfall in 2019, and consequently higher levels of soil moisture, would lead to variation in the strength and direction of selection compared to our original transplant during the historic California drought of 2013. In 2019 absolute levels of soil moisture were 2-3X higher than 2013, and in line with our predictions we observed ultimately higher overall hybrid fitness in 2019 (Figure 3). However the patterns of within-season soil moisture decrease were similar in 2013 and 2019, with moisture decreasing in a slow linear fashion in *M. guttatus’* meadows as opposed to the early season plateau followed by an abrupt and rapid loss of moisture in *M. laciniatus’* rocky habitat. This repeated reciprocal transplant confirms that soil moisture is an important divergent selective factor in *M. laciniatus* and *M. guttatus*’ habitats. Our finding agrees with a recent common garden experiment that also found water availability to be a strong divergent selective force between the closely related species *M. guttatus* and *M. nasutus* (Mantel & Sweigart 2019), as well as studies in other plant systems (Eckhart et al. 2004; Lee & Mitchell-Olds 2013; Dunning et al. 2016). As the climate warms, water availability is predicted to become an increasingly important factor in species survival (Crimmins et al. 2011), especially for plants like *M. laciniatus* which occur in habitats where soil moisture is ephemeral. Understanding how plant species in extreme habitats respond to fluctuations in soil moisture may help us conserve them.

### The Strength of Fruit and Seed Number Correlation Depends Upon the Environment

There are multiple ways to estimate plant lifetime fitness and each measurement has benefits and caveats (Lovett Doust & Lovett Doust 1998). By examining multiple fitness metrics we can learn more about the biology of reproduction and response to selection in our particular system. We have used fruit number as one fitness metric in our phenotypic selection analysis to compare data across years, however in 2019 we also included seed number as a more accurate fitness proxy. Fruit and seed number are significantly correlated in our data, as found in previous research (Baker et al. 1994; Hensen & Wesche 2006), but they do not have the same degree of correlation across habitats. The differences that we see between fruit number and seed number across habitats, with a stronger relationship in meadows, also follow patterns that have been found in other systems. Seeds in flowers produced later on the meristem are commonly aborted due to resource limitation (Stephenson 1981). This is especially common in plants growing in unpredictable environments with truncated growing seasons like granite outcrops. Seed abortion during resource limitation later in the season can produce patterns of weaker seed/fruit correlation like that we see in *M. laciniatus’* granite outcrop environment (reviewed in Lovett Doust & Lovett Doust 1998). In the granite outcrop where water becomes very quickly limited in both years, it is possible that plants abort later fruits and direct most resources to seeds in the first fruit, while in the meadow habitat with extended water availability resources are put towards more flowers and therefore fruits. The differences in the strength of correlation between fruit and seed number between the habitats may explain why selection gradients for flowering time switched direction when seed was used as the fitness metric in 2019. While selection for early flowering was under stronger selection for fruit number than setting any fruit at all, a shift in selection for seed number could mean that plants diverted resources to producing many fruits rather than many seeds in one fruit. In this way seasonal changes in ontogeny can lead to opposing selection gradients. This same pattern of opposing selection gradient signs on the same phenotype has been noted in other reciprocal transplant studies with multiple fitness metrics, including a study in *Clarkias* in which a shift in direction of selection on flowering time occurs from fruit to seed number (Anderson et al. 2015). Because these patterns are not visible in plants grown at optimal conditions in the greenhouse, differences in selection directionality between fruits and seeds may allow us to understand more nuanced differences in resource allocation.

### Temporally Varying Selection Could Erode Species Divergence

Since adaptation to different environments, or habitat isolation, is an important component of reproductive isolation between *M. laciniatus* and *M. guttatus* (Ferris & Willis 2018), we predicted that we would find divergent selection in ecologically important and reproductively isolating traits in the direction of species differences across both time and space. Divergence in flowering time is often important for local adaptation and pre-zygotic reproductive isolation between closely related species (Hall & Willis 2006; Martin et al. 2007; Lee & Mitchell-Olds 2013; Mantel & Sweigart 2019). Earlier flowering time allows for rapid development and reproduction in ephemeral environments (Fox 1989; Kazan & Lyons 2016). Selection for earlier flowering time was stronger in granite outcrops in both years in our truncated poisson model and this matches our predictions since *M. laciniatus* flowers earlier than *M. guttatus*.

However, the wetter 2019 season weakened the strength of divergent selection between habitats and even caused selection in the opposite direction of species differences in our binomial model. These shifts in the strength and direction of selection between years suggest that temporal shifts in the environment may reduce divergent selection between species. A replicated transplant study over dry and wet years in the *Clarkia* system also found that flowering time differentiation persisted over time, with overall earlier flowering in the dry year (Eckhart et al. 2004). More similar flowering time across habitats in wetter years may have implications for hybridization in sympatric populations, potentially eroding species divergence in years with reduced selective pressures (Nagy 1997).

Flower size and stigma-anther separation are also important pre-zygotic isolating barriers, and are associated with the shifts in mating system that we see between our two focal species (Snell & Arson 2005). We predicted to find selection for reduced stigma-anther separation and smaller flower size in the granite outcrop. In 2019 selection gradients in the granite outcrop poisson model indicated directional selection for larger flowers and increased stigma-anther separation, an unexpected result. Previous research has also shown that self-fertilization in *M. guttatus* occurs through pollen transfer on the corolla and the curling of the stigma (Arathi & Kelly 2004), and anther stigma-separation does not have a strong relationship with selfing in *M. guttatus* (Robertson et al. 1994), although this has not been directly tested in *M. laciniatus*. While flowering time has often been found to vary seasonally based on environmental stresses that a plant experiences (Kazan & Lyons 2016), it is not well know in this system how flower size and stigma-anther separation vary based on seasonal environmental stressors. Identifying the importance of extrinsic pre-zygotic barriers, and how they might vary in sympatry versus allopatry, is an important next step in further understanding reproductive isolation and potential reinforcement between the two species (Coyne & Orr 1989, Hopkins 2013). Future research investigating the both sympatric and allopatric populations will allow us to quantify and understand the importance of these barriers. Additionally, while we control for the effects of maternal effects and transgenerational plasticity in our experimental design, we do not quantify how these factors might impact reproductive isolation. Future studies incorporating these factors will further inform our results.

In 2013 we found patterns of selection on leaf lobing in the direction of species differences, selection for lobing in granite outcrops and not meadows. However in 2019 we found selection for leaf lobing based on fruit number in both habitats in our truncated poisson analysis and strong selection against leaf lobing in granite outcrops in the binomial model. Unexpected selection against the direction of species differences (Ferris et al. 2015), has been found in previous studies and may be caused by correlation with other traits (Nagy 1997). Flowering time is under selection in the same direction in both sites, and flowering time and leaf lobing are negatively correlated in the meadow habitat. Leaf lobing could also be correlated with an unmeasured phenotype. Previous QTL mapping has shown that leaf lobing is a complex trait controlled by three large effect QTLs (Ferris et al. 2015), with the largest effect QTL influencing both leaf lobing and flowering time (Ferris et al. 2017). Under this QTL is candidate locus TCP4 which has been found in tomatoes (Ori et al. 2007) and *Arabidopsis thaliana* to play a large role in both leaf cell differentiation and floral development (Nag et al. 2009). While the phenotypic selection models take into account the effect of other traits in determining selection gradients, linkage and pleiotropy may also play an important role in determining patterns in leaf lobing and flowering time across populations.

### Conclusion

We found that temporally and spatially varying selection affect species divergence in two sympatric *Mimulus* species by repeating a a previously published reciprocal transplant (Ferris and Willis 2018) experiment six years later and measuring selection and traits in advanced generation hybrids. Soil moisture was closely tied to fitness across space and time and in a year with increased soil moisture we saw a decrease in the strength of selection on reproductive isolating and locally adaptive traits. Our findings that spatially varying selection is a stronger force than temporally varying selection in maintaining species differences supports previous findings in other systems (reviewed in Kassen 2002 and Siepielski et al. 2013). Kassen (2002) found that in heterogeneous environments spatially varying selection often led to the evolution of specialists, while temporally varying selection led to the evolution of generalists. In reference to our results, this may suggest that continued temporal fluctuations in environmental conditions could cause divergent species to become more similar over time. A long-term study of Galapagos finches also found that temporal fluctuations in selection and severe drought conditions resulted in closely related species become more similar through increased introgression (Grant & Grant 2020). While spatial and temporal selection may be comparable in magnitude and variation, this relationship is confounded by interactions of spatial and temporal selection that are not accounted for in the published literature (Siepielski et al. 2013). Our synchronous measurement of spatial and temporal selection allows us to disentangle the two forces and investigate their importance in maintaining species boundaries.

Repeated studies such as ours are essential for a better understanding of how spatial and temporal shifts in environmental factors alter selective pressure on quantitative trait divergence between species, including traits contributing to reproductive isolation. Repeated reciprocal transplant experiments have been instrumental in identifying local adaptation in both individual species (Waser & Price 1985; Eckhart et al. 2004; Agren & Schemske 2012; Anderson et al. 2021) and hybrid zones (Wang et al. 1997), but ours is one of the few to identify how both spatial and temporal variation in selection contributes to or erodes species divergence. While divergence and speciation are often viewed as directional processes, especially in reference to self-fertilizing species that have lost much of the genetic variation that precedes differentiation (Stebbins 1957), it is important to acknowledge the more nuanced and varying nature of species boundaries. Our research suggests that relaxed selective pressures in a wetter year erodes reproductive isolation and species divergence. However, given the effects of climate change in California it is likely that most future years will be dry, resulting in stronger divergent selection similar to our 2013 year transplant. While the continued warming and drying of our planet is a cause of great concern for the continuance of many species, in this specific case it may strengthen species boundaries through powerful micro-habitat mediated selection in favor of local genotypes. The implications of rare wet years with relaxed selection in longer term patterns of species’ divergence remains to be further understood. Continued studies of simultaneous spatial and temporal variation in selection will allow us to better understand how shifting environmental conditions shape species boundaries and persistence in a changing world.

## Acknowledgments

We would like to acknowledge and pay tribute to the original inhabitants of the unceded land on which our research was conducted, the Southern Sierra Miwuk Nation, Bishop Paiute Tribe, Bridgeport Indian Colony, Mono Lake Kutzadikaa, North Fork Rancheria of Mono Indians of California, Picayune Rancheria of the Chukchansi Indians, and the Tuolumne Band of Me-Wuk Indians. Their culture and stewardship remain an integral part of the land. As scientists we strive to take responsibility for the impacts of colonialism in our field and move forward with respect and support of indigenous movements and knowledge. Thank you to Yosemite National Park Service for logistical, community, and permitting support. This work was performed (in part) at the University of California Natural Reserve System Yosemite Reserve DOI: 10.21973/N3V36C. Many thanks to the Blackman Lab at UC Berkeley and Sexton Lab at UC Merced for providing greenhouse and growth chamber resources. Thank you to Kasandra Scholz, Bridget Dixon, and Kathleen Sway for their work measuring leaf shape and counting seeds. We are grateful to two anonymous reviewers for their constructive and thoughtful comments.

## End Section Statements

### Ethics

Field work was conducted under YNP Permit #YOSE-00831.

### Data, code, and material

Data and code available from the Dryad data repository https://doi.org/10.5061/dryad.prr4xgxpk.

### Author Contributions

DT collected field data, carried out statistical analysis, and drafted the manuscript; ECW collected field data; KGF conceived of the study, designed the study, provided funding, assisted in field work and statistical analysis, and critically revised the manuscript. All authors gave final approval for publication and agreed to be held accountable for the work performed therein.

## Supplementary Data

**Figure S1.**
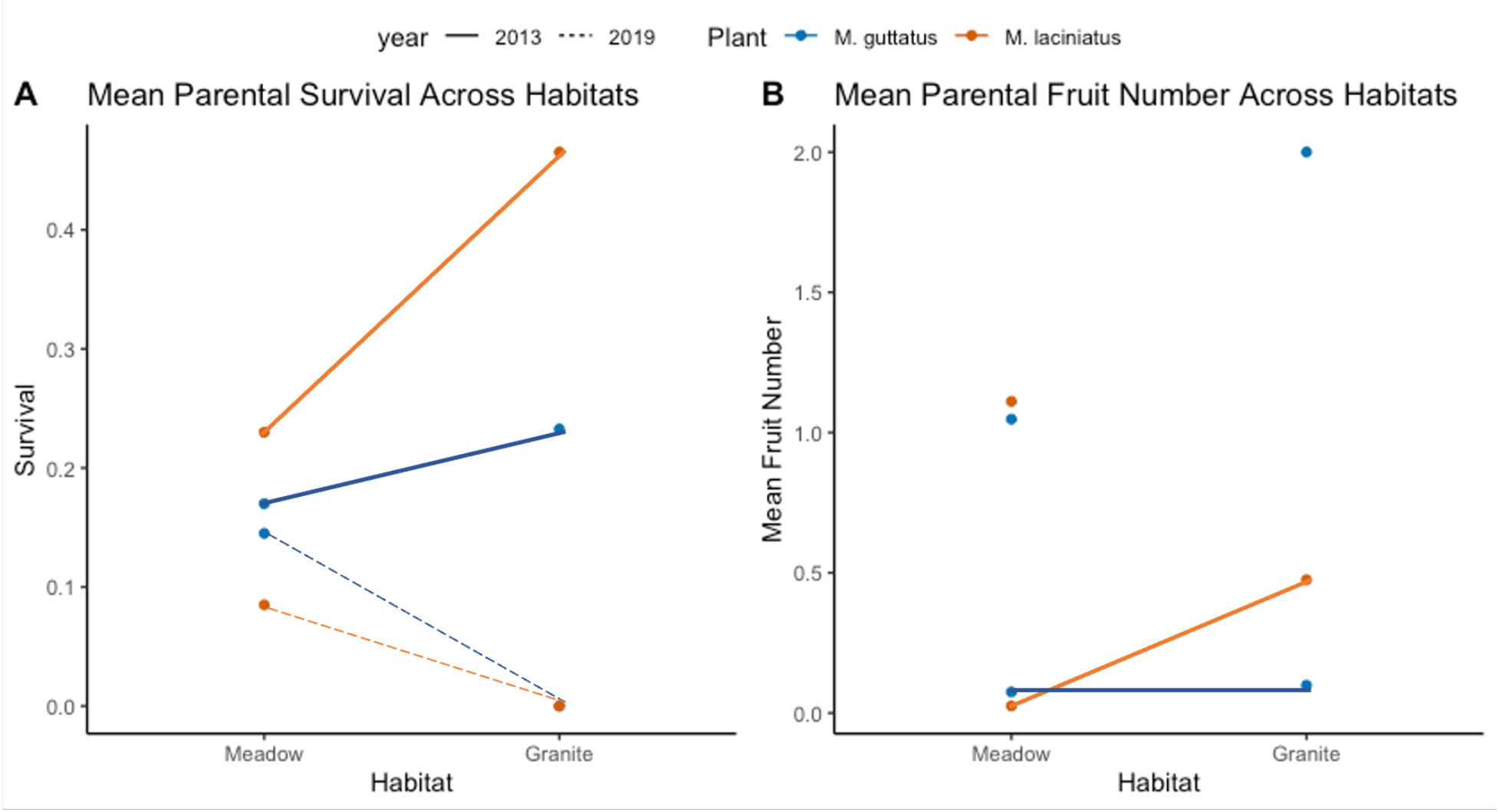
A) Mean parental species survival across year and habitat. B) Mean parental fruit number across habitats. Lines indicating relationships across habitats are not included for 2019 because sample sizes are insufficient. 2013 data is published in Ferris and Willis (2018) and re-analyzed here.

**Figure S2.**
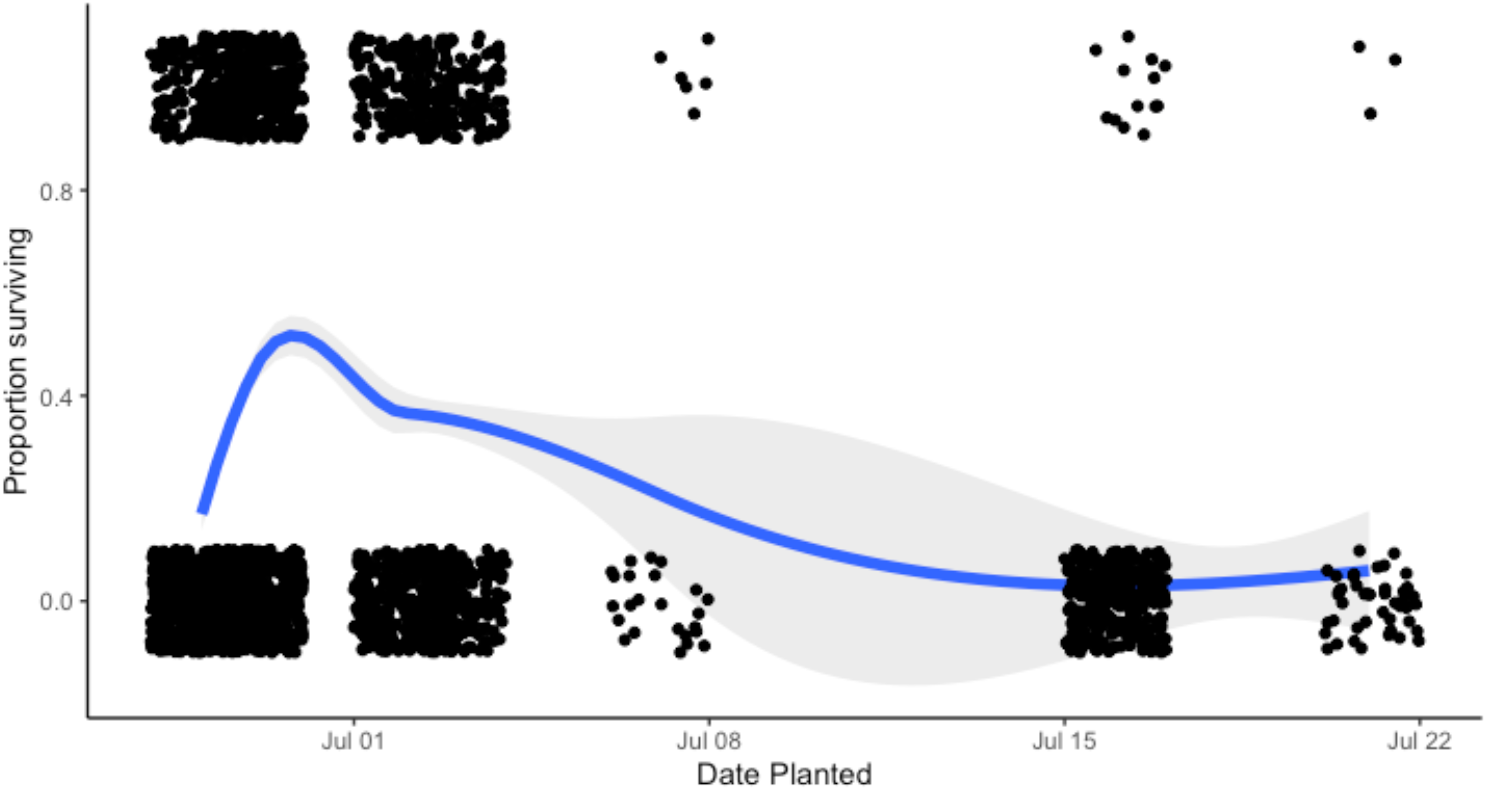
Proportion of individuals surviving in all sites based on planting date. The line indicates a summary line with a 95% confidence interval. A one-way ANOVA indicates that there is a significant difference in plant survival based on planting date (F=77.2, p<.001).

**Figure S3.**
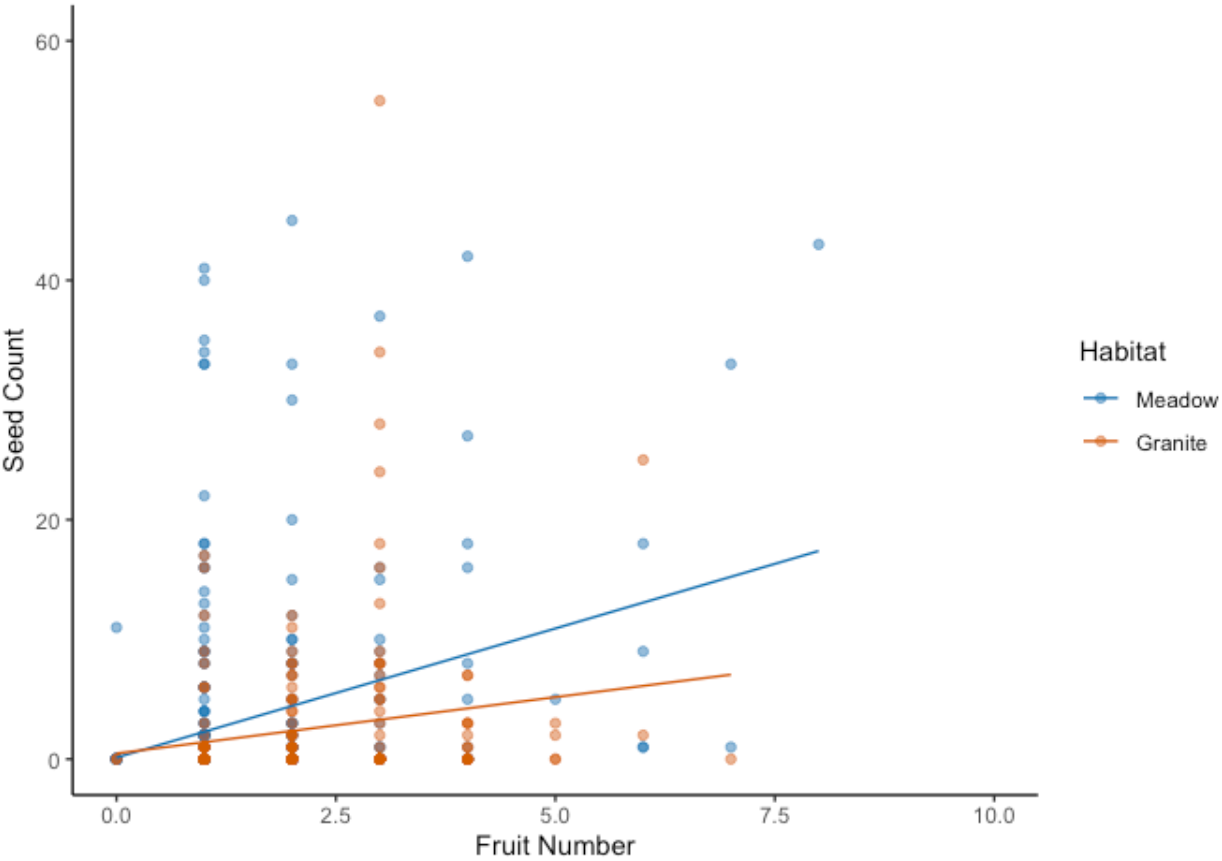
Correlation between seed and fruit is significant in meadows (P<.001, correlation coefficient = 0.309, n = 1902) and the granite outcrop (P=0.0036, correlation coefficient = 0.172, n=838).

**Figure S4.**
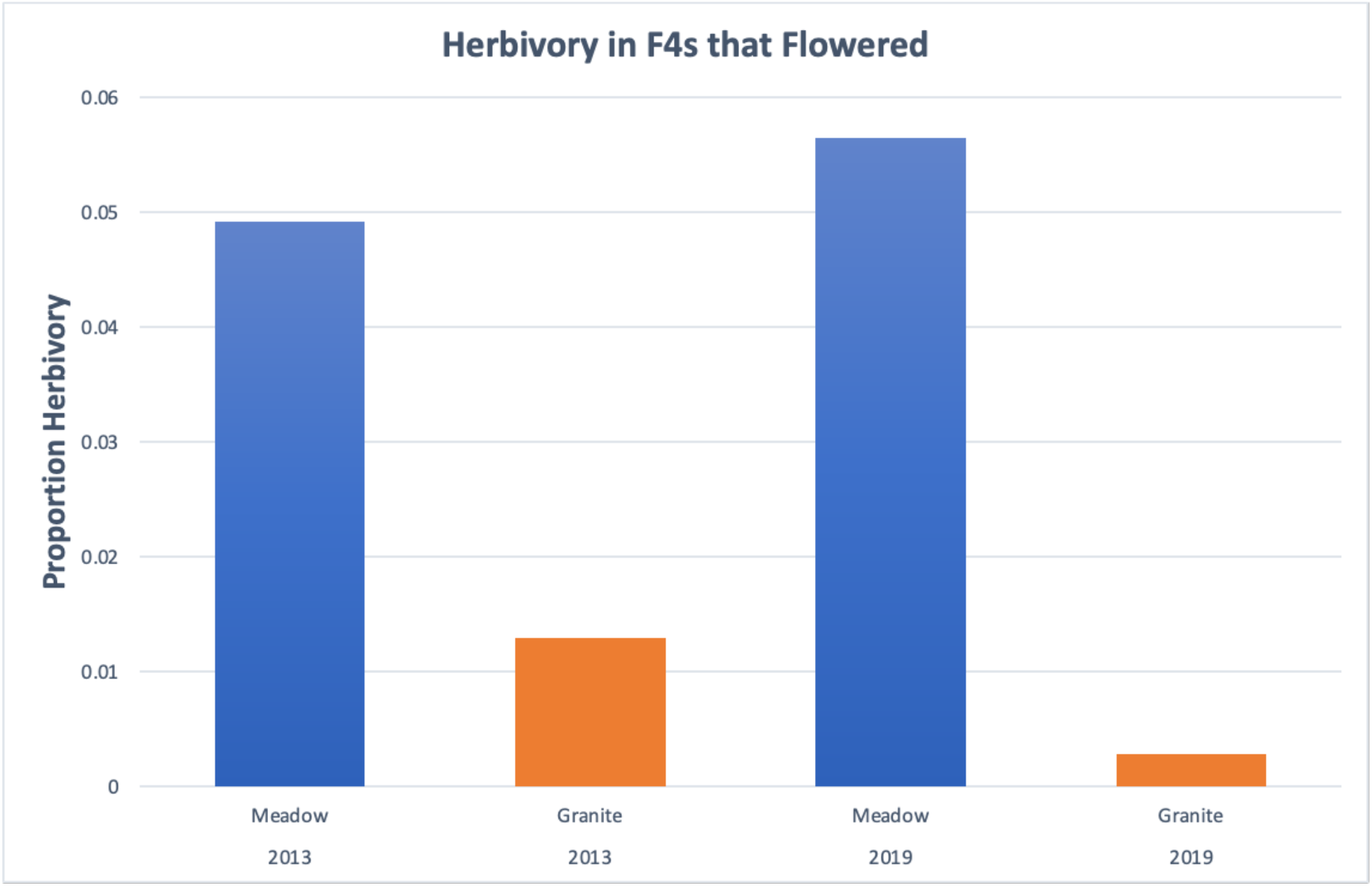
Proportion of total F4 hybrid individuals that flowered with presence of herbivory in 2013 and 2019. 2013 data is published in Ferris and Willis (2018) and re-analyzed here.

**Figure S5.**
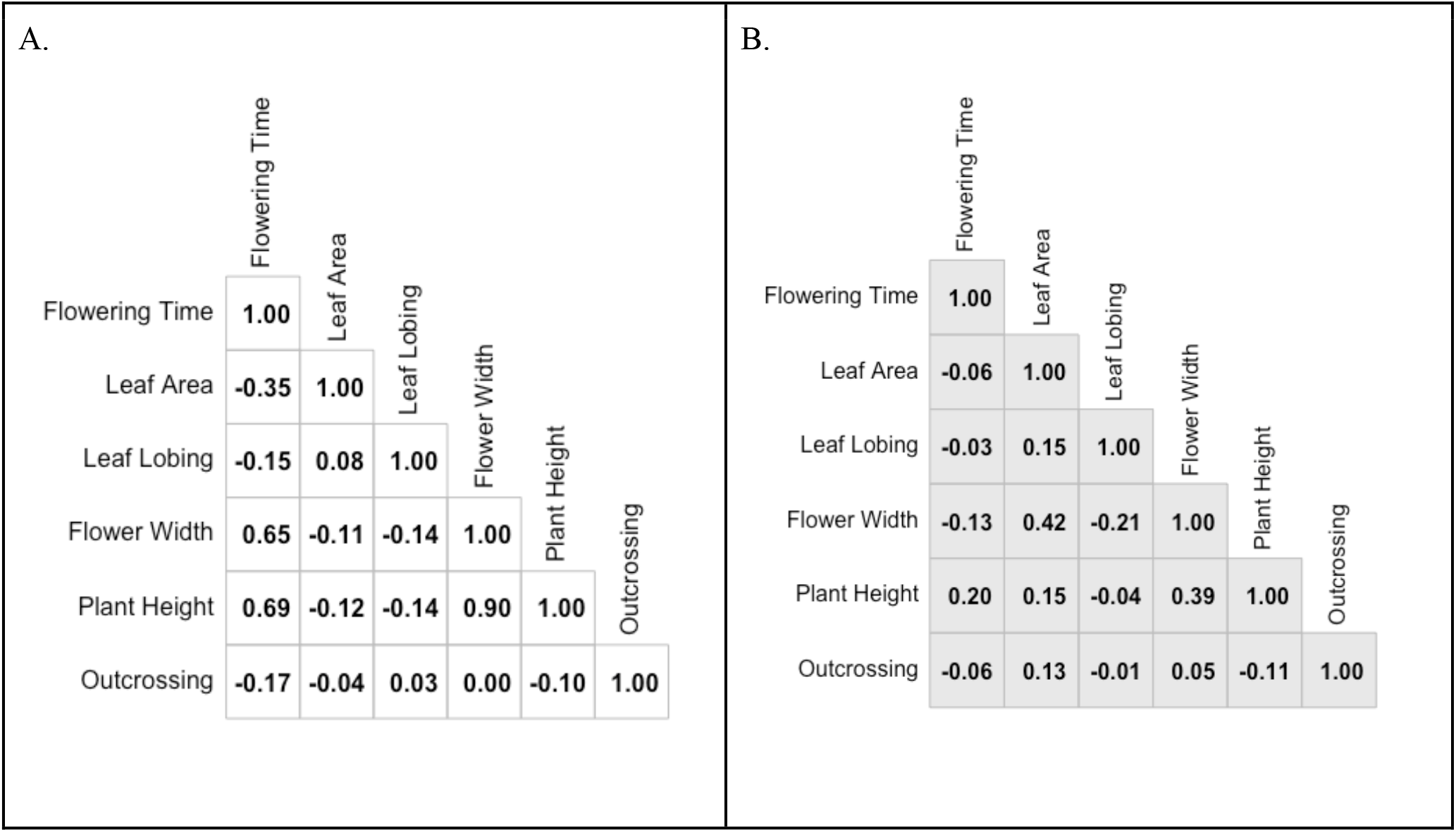
Trait correlation matrix for F_4_ hybrid traits in 2019 in A.) Meadow and B.) Granite.

**Figure S6.**
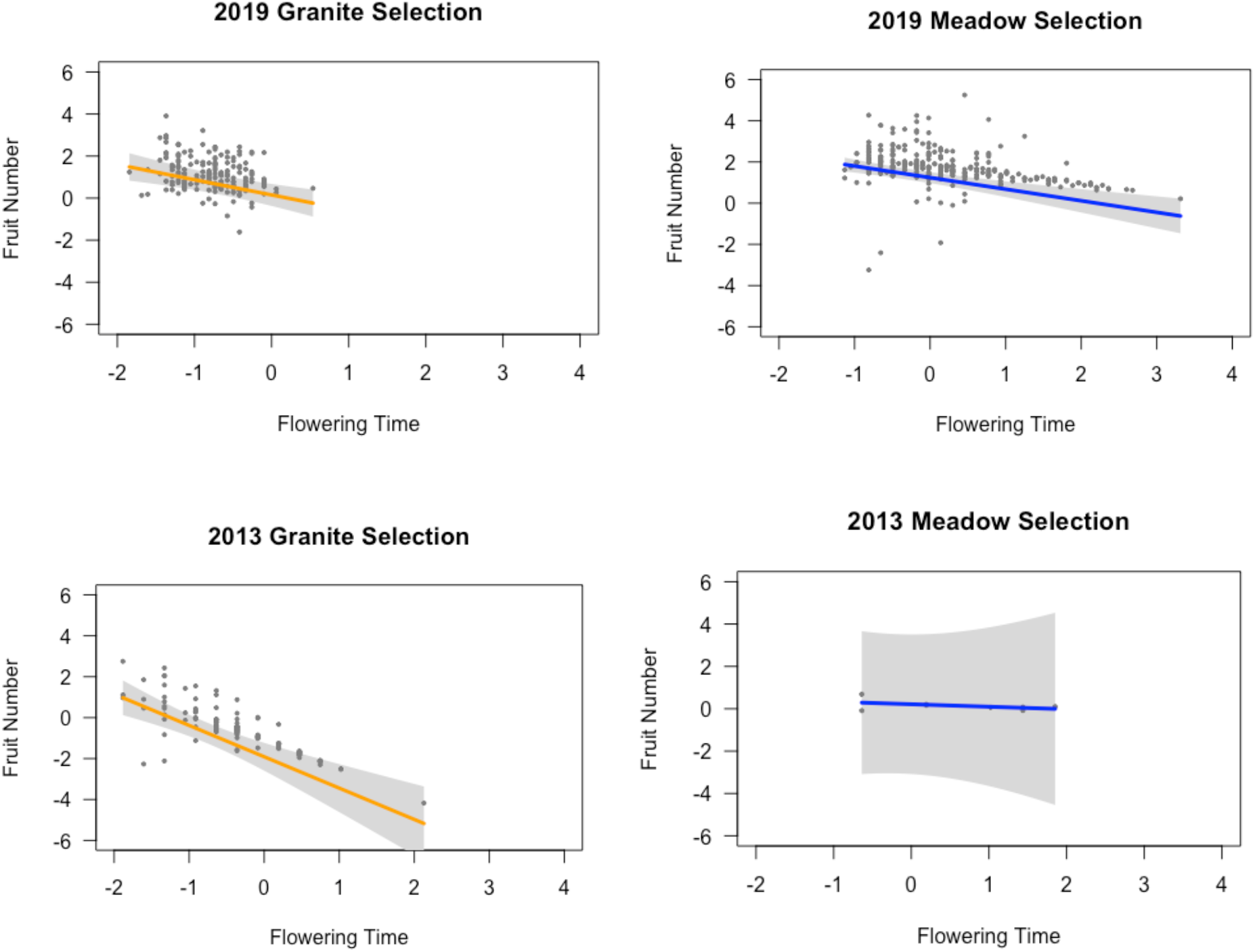
Partial residuals from the multiple regressions in the truncated poisson models showing selection on flowering time while holding all other explanatory models at their median value for granite habitats (orange) and meadow habitats (blue). Models are visualized using the R package visreg (Breheny & Burchett 2017). 2013 data is published in Ferris and Willis (2018) and re-analyzed here.

**Figure S7.**
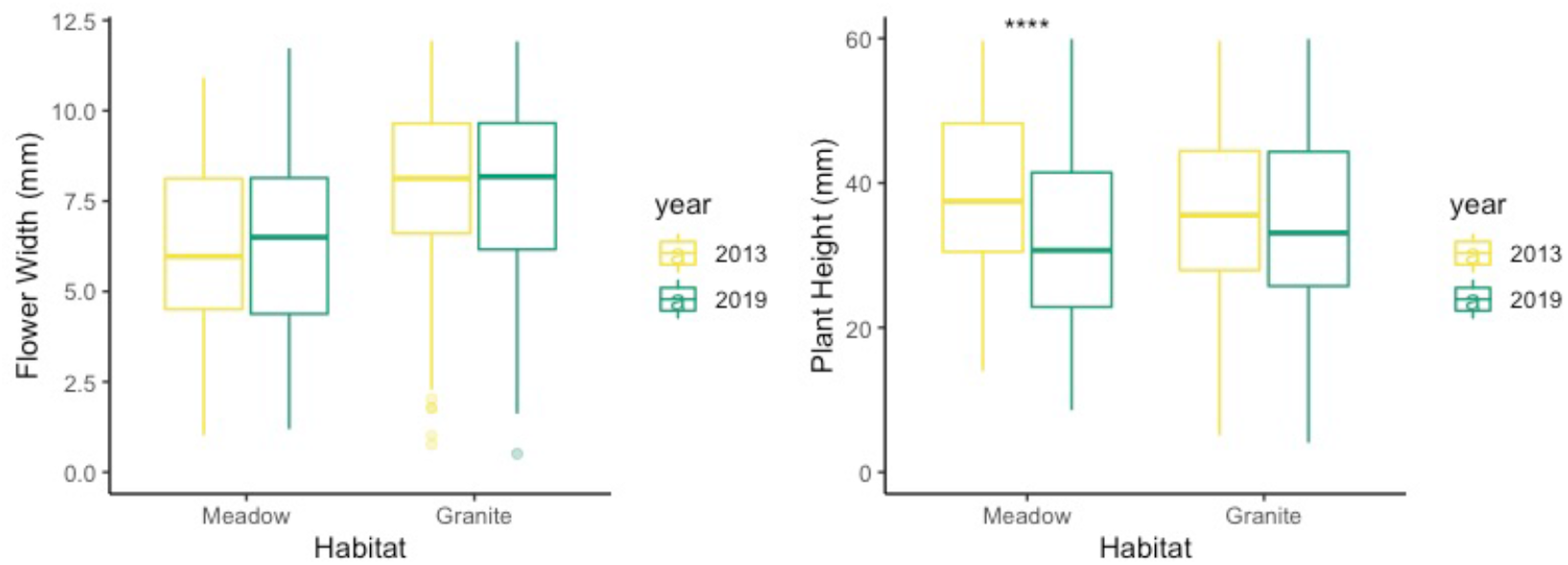
Flower width and plant height in F4 hybrids (base to meristem) across years and habitats. Yellow boxes are 2013 data and green boxes are 2019 data. Significance difference between year in a trait is indicated by “*” and significance codes are as follows: P-value < 0.0001****; 0.001***; 0.01**; 0.05*. 2013 data is published in Ferris and Willis (2018) and re-analyzed here.

**Table S1.** Model selection for best fit models of plant survival and environmental variables. For each analysis, we report k, the number of parameters; Log(L), the likelihood score; AIC score, delta(i), the difference between that model’s AIC and the top model’s AIC; and wi, the model weight.

| Year | Model | k | Log(L) | AIC | delta(i) | wi |
| --- | --- | --- | --- | --- | --- | --- |
| 2019 | Survival Total ~ moisture*days*Habitat | 11 | 272.649 | -522.9 | 0 | 1 |
| 2013 | Survival Total ~ moisture*days*habitat | 11 | 488.0297 | -954.06 | 0 | 0.499 |
| Both | Survival Total ~ moisture*days+Habitat+year | 9 | 318.120 | -618.2 | 0 | 0.88 |
| Both | Survival Meadow ~ moisture+days*Site+year | 8 | 395.961 | -775.8 | 0 | 0.958 |
| Both | Survival Granite ~ moisture*days+Site+year | 8 | 236.335 | -456.6 | 0 | 1 |

**Table S2.**
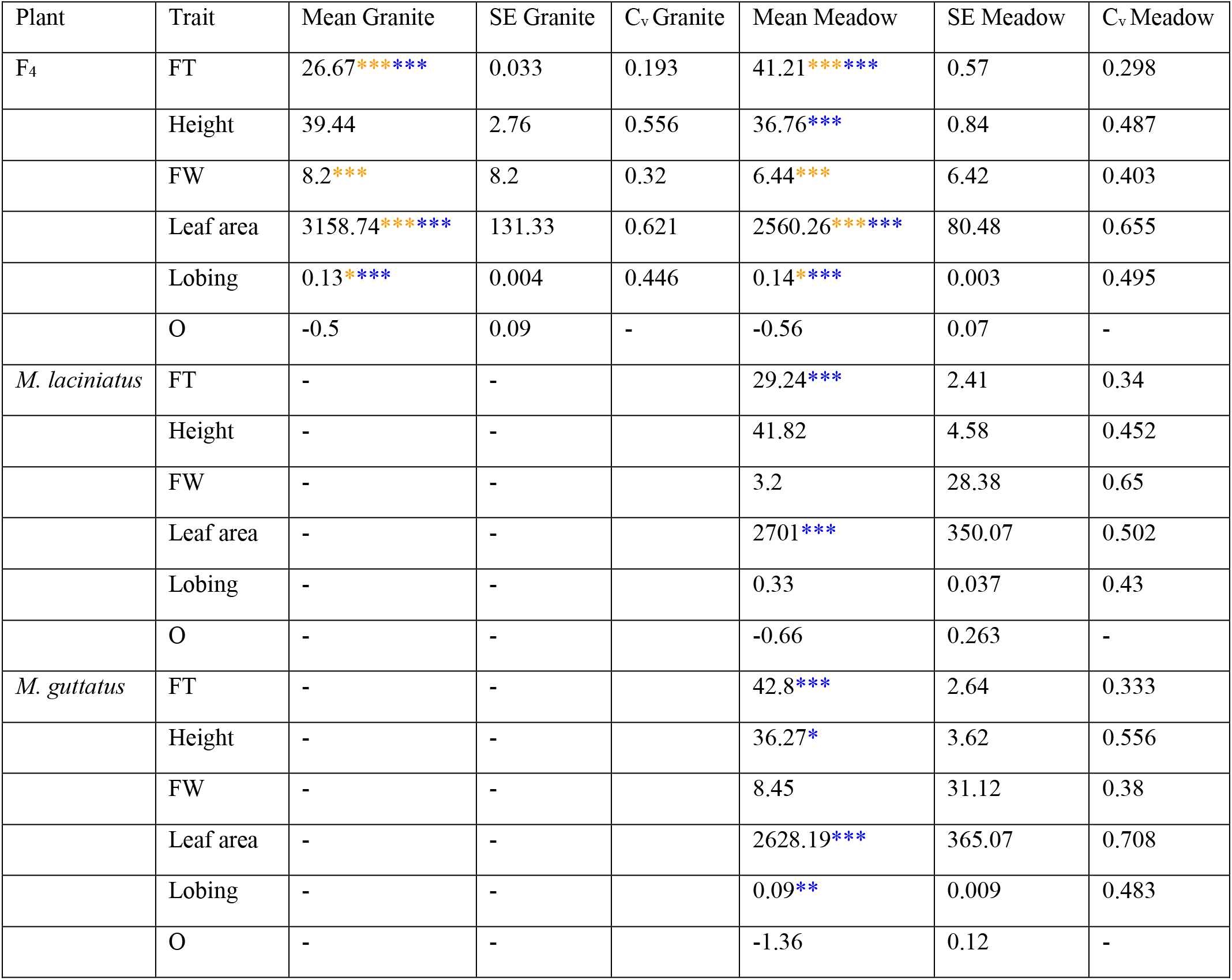
Trait means, standard error, and coefficient of variance (C_v_) from the 2019 experiment. Significant variation in a trait mean is indicated by “*” within year and “*”across years, and significance codes are as follows: P-value < 0.001***; 0.01**; 0.05*. FT, flowering time; FW, corolla width; O, for stigma-anther separation.

**Table S3.** Model selection for best fit linear and quadratic selection models of hybrid traits in 2019. For each analysis, we report k, the number of parameters; Log(L), the likelihood score; AIC score, delta(i), the difference between that model’s AIC and the top model’s AIC; and wi, the model weight. FT, flowering time; FW, corolla width; L, leaf lobing; PH, height; O, for stigma-anther separation.

| Model | k | Log(L) | AIC | delta(i) | wi |
| --- | --- | --- | --- | --- | --- |
| Binomial |  |  |  |  |  |
| Granite Fruit ~ FT X FW + L X PH | 10 | -81.038 | 183.4 | 0 | 0.999 |
| Granite Seed ~ FT | 8 | -99.608 | 216.1 | 0 | 0.334 |
| Meadow Fruit ~ FT X FW + L X PH | 10 | -145.679 | 312.1 | 0 | 0.195 |
| Meadow Seed ~ FT X PH + FW | 8 | -162.887 | 342.3 | 0 | 0.431 |
| Truncated Poisson |  |  |  |  |  |
| Granite Fruit ~ L X PH + FW X FT | 9 | -224.734 | 467.5 | 0 | 0.445 |
| Granite Seed ~ FT X O + PH X FW | 9 | -222.483 | 466.4 | 0 | 0.947 |
| Meadow Fruit ~ L X PH + FT | 7 | -257.548 | 529.5 | 0 | 0.241 |
| Meadow Seed ~ FT X PH + FW X L | 9 | -1012.73 | 2045.2 | 0 | 0.999 |

**Table S4.** Model selection for best fit linear and quadratic selection models of hybrid traits in 2019 including planting time. For each analysis, we report k, the number of parameters; Log(L), the likelihood score; AIC score, delta(i), the difference between that model’s AIC and the top model’s AIC; and wi, the model weight. FT, flowering time; FW, corolla width; L, leaf lobing; PH, height; O, stigma-anther separation; PT, planting time.

| Model | k | Log(L) | AIC | delta(i) | wi |
| --- | --- | --- | --- | --- | --- |
| Binomial |  |  |  |  |  |
| Granite Fruit ~ FT X FW X PT + L X PH | 14 | -72.775 | 176.1 | 0 | 0.53 |
| Granite Seed ~ FT | 8 | -99.608 | 216.1 | 0 | 0.334 |
| Meadow Fruit ~ FT X FW + L X PH X PT | 14 | -138.765 | 307.0 | 0 | 0.417 |
| Meadow Seed ~ FT X PH + FW | 8 | -162.887 | 342.3 | 0 | 0.431 |
| Truncated Poisson |  |  |  |  |  |
| Granite Fruit ~ L X PH X PT + FW X FT | 13 | -215.95 | 460.2 | 0 | 0.663 |
| Granite Seed ~ FT X O + PH X FW | 9 | -222.483 | 466.4 | 0 | 0.947 |
| Meadow Fruit ~ L X PH + FT + PT | 8 | -254.295 | 525.1 | 0 | 0.425 |
| Meadow Seed ~ FT X PH + FW X L | 9 | -1012.73 | 2045.2 | 0 | 0.999 |

